# RNA-Seq Time Series of *Vitis vinifera* Bud Development Reveals Correlation of Expression Patterns with the Local Temperature Profile

**DOI:** 10.1101/2020.10.18.344176

**Authors:** Boas Pucker, Anna Schwandner, Sarah Becker, Ludger Hausmann, Prisca Viehöver, Reinhard Töpfer, Bernd Weisshaar, Daniela Holtgräwe

## Abstract

Plants display sophisticated mechanisms to tolerate challenging environmental conditions and need to manage their ontogenesis in parallel. Here, we set out to generate an RNA-Seq time series dataset throughout grapevine (*Vitis vinifera*) early bud development. The expression of the developmental regulator *VviAP1* served as an indicator for progress of development. We investigated the impact of changing temperatures on gene expression levels during the time series and detected a correlation between increased temperatures and a high expression level of genes encoding heat-shock proteins. The data set also allowed the exemplary investigation of expression patterns of genes from three transcription factor (TF) gene families, namely MADS-box, WRKY, and R2R3-MYB genes. Inspection of the expression profiles from all three TF gene families indicated that a switch in the developmental program takes place in July which coincides with increased expression of the bud dormancy marker gene *VviDRM1*.

## 1. Introduction

Plants are sessile organisms which cannot escape from herbivores or changes in environmental conditions. As a consequence, various stress response mechanisms [1,2] and complex specialized metabolite pathways evolved to counteract adverse situations and conditions [3,4]. These response mechanisms also need to protect plant embryogenesis, as well as vegetative developmental processes like outgrowth of side shoots from resting or latent buds. All these developmental processes, including the establishment of compound buds, have to proceed undisturbed despite potentially challenging and/or unfavorable environmental conditions.

Similar to other woody perennial plants like e.g. apple or poplar, *V. vinifera* (grapevine) bud development spans over two years between bud initiation and growth of new side shoots. Newly formed buds enter in a dormancy phase in the winter time between the two growing seasons before buds sprout in the second season [5,6]. In spring of the first season (April/May on the northern hemisphere), new axillary buds are formed on young grapevine shoots. These new buds initially contain meristems that develop into embryonic shoots with their shoot apical meristems (SAM) and containing primordia for leaves, tendrils and inflorescences. This implies that different types of meristems, including lateral and inflorescence meristems, co-exist in the buds. Floral transition takes place at about June, five to seven weeks after burst of “old” buds (i. e. the buds that are one year ahead in development). Inflorescence primordia differentiate from uncommitted primordia formed within the new buds. Due to further differentiation of inflorescence meristems into inflorescence branch meristems (about July), the compound buds finally contain the embryonic version of next year’s shoots, each with tissues for first leaves, inflorescences and tendrils [6,7]. The buds enter endodormancy which passes over into ecodormancy depending on the environmental conditions of fall and winter [8–10]. In early spring of the second season, ecodormancy is released and inflorescence branch meristems produce single flower meristems in swelling buds (April) and flower organ development begins [6]. It is important to note that the precise timing of floral transition and development strongly depend on environmental conditions and genotype.

Heat-shock proteins (HSPs) are a group of proteins, which were initially detected due to their accumulation in response to quickly increased temperature. They realize a molecular mechanism to endure higher temperatures. First reports of HSPs in plants reach back to the 1980s when they were described based on cell culture experiments with tobacco and soybean [11]. HSPs are assumed to support several physiological functions under normal growth conditions. This includes folding, unfolding, localization, accumulation and degradation of other proteins [12,13]. Additionally, irreversible aggregation of other proteins is prevented and refolding is facilitated under heat stress [14]. Several categories of HSPs based on sequence homology and typical molecular weight have been defined [12], thus leading to multiple polyphyletic groups of HSPs.

WRKY transcription factors (TFs) are a family of TFs, which play an important role in the regulation of responses to environmental stress conditions [15–17]. R2R3-MYB TFs are often responsible for controlling the formation of specialized metabolites in response to environmental triggers, but also regulate several plant-specific processes including root hair and trichome differentiation [18–20]. MADS-box TFs are typically involved in the regulation of developmental processes like determination of plant organ identity [21–23]. One especially important developmental regulator is APETALA1 (AP1), also a MADS-box factor, which connects signals received from the environment with initiation and/or progress of developmental processes [24,25]. *VviAP1* and *VviAIL2*, a *V. vinifera* homolog of the MADS-box gene *AINTEGUMENTA-like* (*AtAIL1*, At1g72570), have been postulated to be involved in the photoperiodic control of seasonal growth [26]. In addition, marker genes for the dormant state of buds have been described. One such marker gene is *DRM1*, a gene that has been found initially in *Pisum sativum* to encode a dormancy-associated protein [27]. Subsequently, *DRM1* homologs have been identified in many species in the context of bud dormancy, including *V. vinifera* [10,28].

In the model plant *Arabidopsis thaliana*, phylotranscriptomic evidence for a molecular embryonic hourglass was published [29–31]. We attempted to create an RNA-Seq dataset to examine early bud development of *V. vinifera* for a similar general pattern. While an hourglass pattern was not detected in the data (M. Quint, personal communication), we harnessed the time series of *V. vinifera* RNA-Seq samples to investigate changes in gene expression during early bud development throughout the first season at a fine scale and observed a strong influence of high temperatures on the expression of HSP genes of field grown plants. In addition, a switch in the expression patterns of various TF genes was observed that happens in parallel to or shortly after the switch from uncommitted primordia to inflorescence primordia. This switch in expression pattern coincides with onset of expression of the dormancy marker gene *VviDRM1*.

## 2. Results

### 2.1. RNA-Seq Time Series of Early Bud Development and Transcript Accumulation Patterns of Selected Marker Genes

Young buds of *V. vinifera* ‘Calardis Musqué’ were harvested in a vineyard in the south of Germany over a period of 156 days of the first season of development, covering the time from June 1st to November 3rd of 2016 (Figure 1) (File S1). Per time point, buds derived from three vines were harvested and subjected to RNA-Seq analyses in triplicates per time point. Values for gene expression, inferred from values for transcript accumulation, were calculated for all transcribed genes. (File S2, File S3, File S4; see methods for details). Time points with only two successful biological replicates were included in the submission to the European Nucleotide Archive (ENA) database (File S1), but excluded from the investigations presented here.

**Figure 1.**
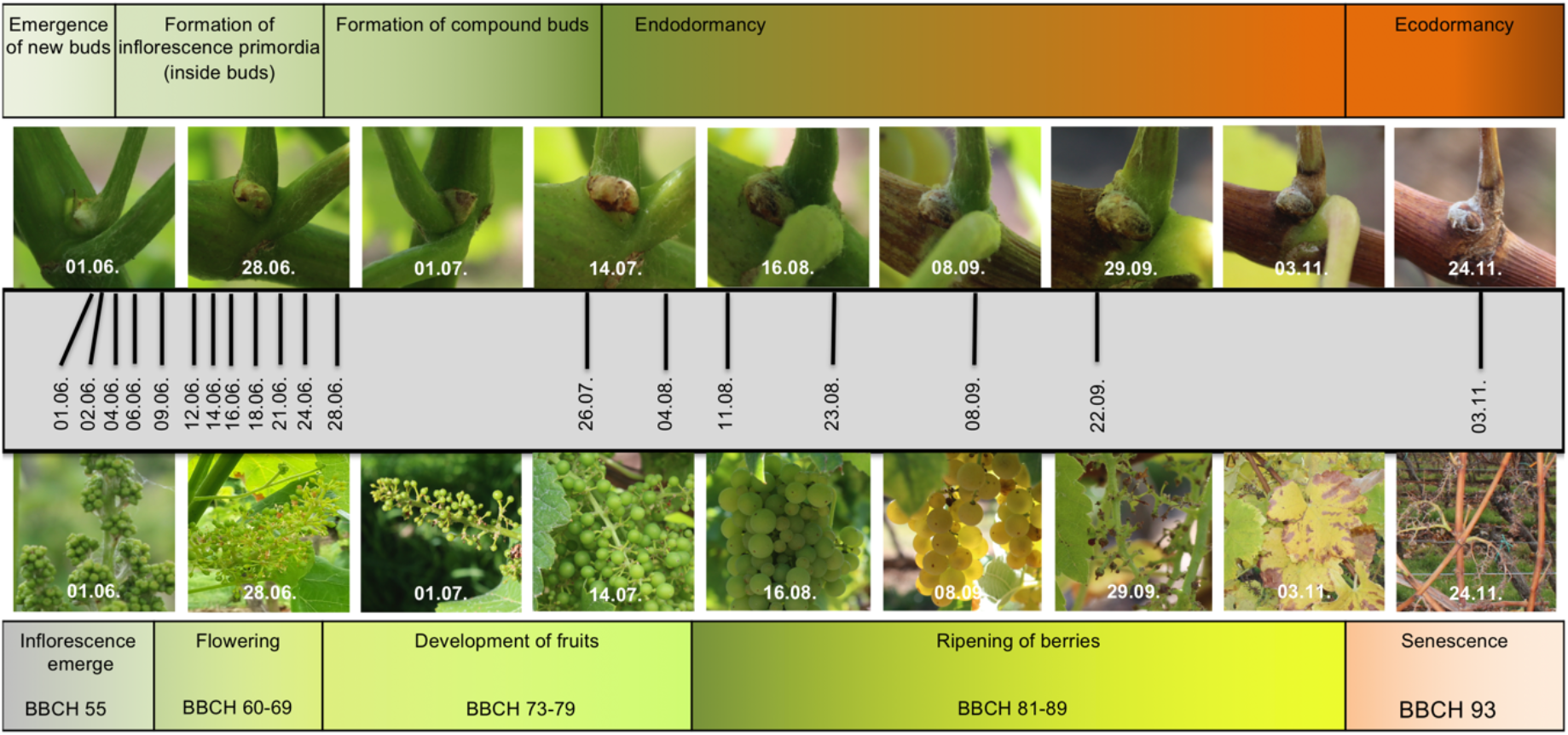
Documentation of the grapevine material used for sampling. The upper row of pictures displays the bud stages during the first year of development, which were the target of this study. The lower row of pictures displays the growth status of the vines from which the young buds were taken. In the center (grey bar), the sampling time line is depicted.

The expression pattern ofthe MADS-Box genes *VviAP1* (CRIBI2.1 ID VIT_201s0011g00100), *VviAIL2* (VIT_209s0002g01370), and *VviSOC1a* (VIT_215s0048g01250) are displayed in Figure 2a. *VviAP1* transcript levels were zero or very low until end of June and rise until September. Due to the time distance between the sampling points we interpret the data as essentially one peak in September. *VviAP1* and *VviAIL2* display quite similar expression patterns. The increase of *VviAP1* transcript levels at the end of June correlates with the time when floral transition, the differentiation of uncommitted primordia into inflorescence primordia, took place. There is no direct correlation between the transcript levels of *VviAP1* and *VviAIL2* with the day length, but the rise of transcript levels coincides with the beginning of reduction of day length after midsummer. We also checked the expression of the dormancy marker gene *VviDRM1* (VIT_210s0003g00090) and found high transcript levels of this gene in the buds with a clear increase starting in July (Figure 2b).

**Figure 2:**
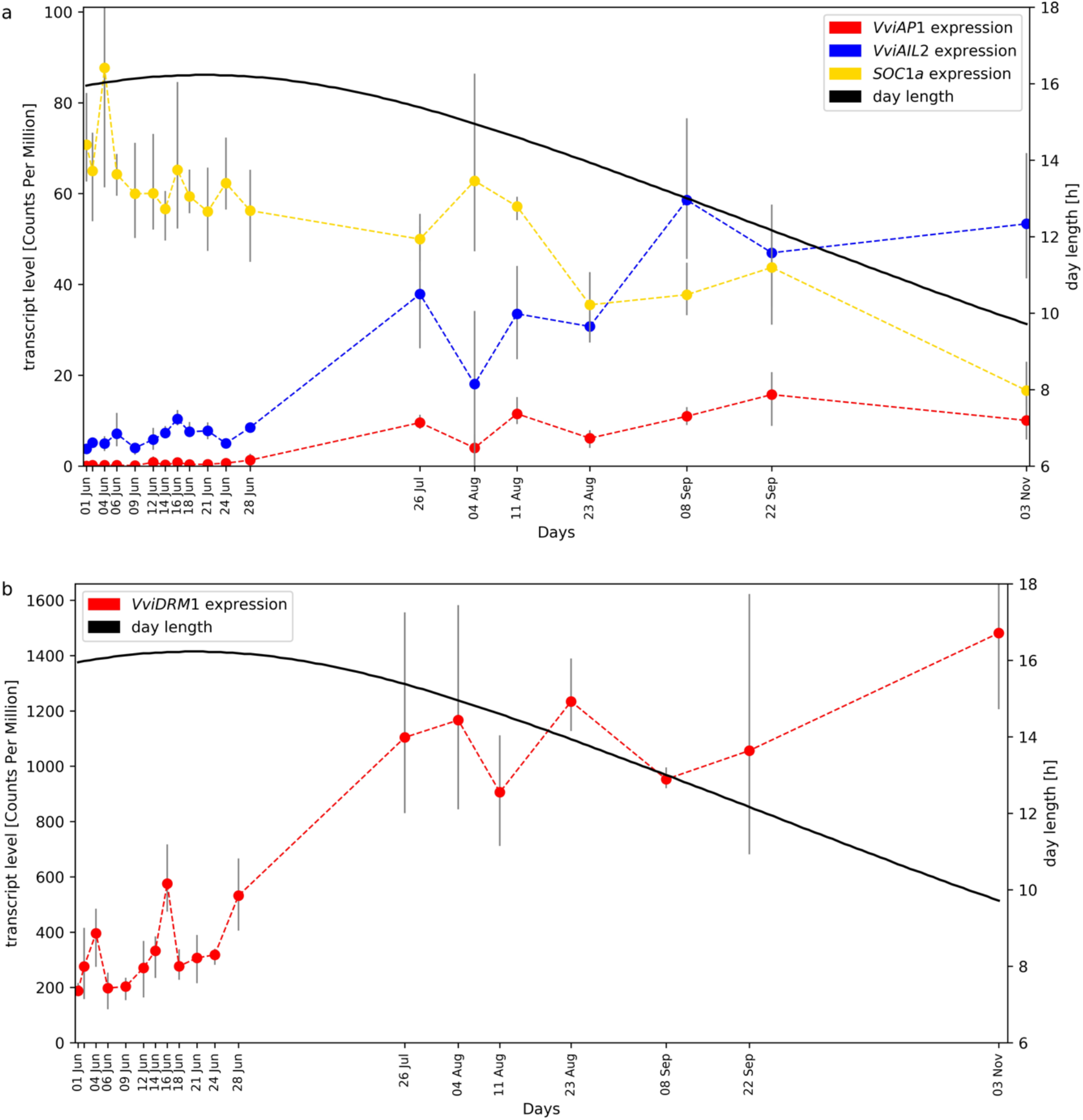
Gene expression time course of *VviAP1, VviAIL2,* and *SOC1a* (panel a) as well as *VviDRM1* (panel b) in developing grapevine buds. The plots were separated due to the large difference in transcript levels (detected as standardized read counts), see y-axis on the left. Day length during the sampling interval is plotted in both panels.

### 2.2. Average Gene Expression Values of HSP Genes Reflect the Local Temperature Profile

We made use of the weather data recorded at the vineyard from which the samples for RNA-Seq were derived. Since the recorded temperature profile during 2016 displayed significant oscillation, we tested the hypothesis that a heat-shock response may take place in the buds. A total of 131 putative HSP genes were identified based on the annotation (File S5), but only 80 of these show substantial transcript abundance (CPM value of more than 10). The gene expression pattern of this set of 80 HSP genes shows a good correlation (r≈0.45) with the temperature profile. Obviously, the correlation is much better during the period with dense sampling. We observed clear HSP gene transcript level peaks at time points with high temperatures (Figure 3). This is especially noticeable at June 24th when the highest average temperature was recorded. A correlation of the day length/photoperiod with the expression pattern of these genes was not observed.

**Figure 3:**
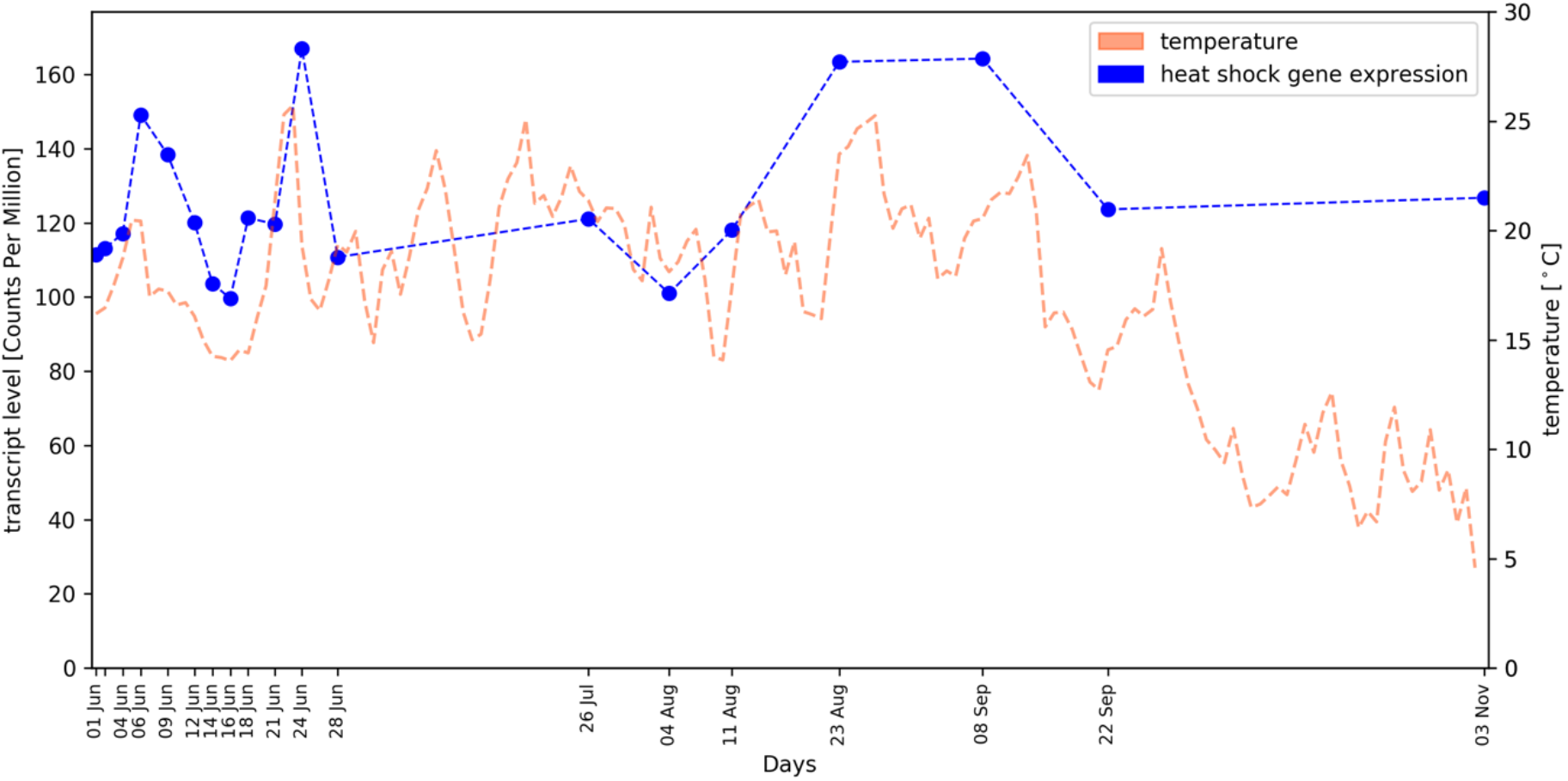
Correlation of HSP gene expression (blue dots / bue dotted line) in developing grapevine buds and environmental air temperature (red dashed line); the course of of the daily average temperature values is shown.

### 2.3. Investigation of Transcription Factor Gene Families: WKRY, MADS-box, and R2R3-MYBs

We harnessed the presented RNA-Seq time series for the analyses of expression patterns of three TF gene families. Heatmaps display transcript levels of genes encoding MADS-box (File S6), R2R3-MYB (File S7), and WRKY (File S8) TFs. Only genes that display detectable transcript accumulation values were considered (see Methods for the threshold). As can be seen from all three heatmaps, the gene activity patterns of quite some of the transcription factor genes change quite dramatically with the onset of *VviAP1* transcript accumulation between June 28th and July 26th.

While *VviSVP1* (VIT_200s0313g00070), an ortholog of the *A. thaliana* MADS-box gene *SHORT VEGETATIVE PHASE*/*AGL22* (At2g22540), shows transcript levels with almost constant values, the gene *VviFLC2* (VIT_214s0068g01800) displays a time course quite similar to those of *VviAP1* and *VviAIL2*. *VviFLC2* is, like its paralog *VviFLC1* (VIT_201s0010g03890), closely related to the *A. thaliana* MADS box gene *FLOWERING LOCUS C*/*AGL25* (At5g10140) which encodes a central repressor of the floral transition. In contrast, *VviTM8a* (VIT_217s0000g01230), which was named according to a gene initially detected in *Solanum lycopersicum* (*TOMATO MADS 8*) that became “founder” of a specific sub-clade of evolutionary related MADS-box genes, shows a transcript accumulation peak at the end of June. Finally, *VviSOC1a* (VIT_215s0048g01250), a homolog of the *A. thaliana* MADS box gene *SUPPRESSOR OF OVEREXPRESSION OF CO 1*/*AGL20* (At2g45660) displays transcript accumulation values that decline after middle of August (Figure 2a).

With regard to the R2R3-MYB genes, *VviMYBC2-L1* (VIT_201s0011g04760), *VviMYB4A* (VIT_203s0038g02310), and *VviMYBPAR* (VIT_211s0016g01300) are prominent examples of genes that support the gene expression pattern switch during July. In addition to the July switch, several *R2R3-MYB* genes display high transcript accumulation values specifically in November. Clear examples are *VviMYB15* (VIT_205s0049g01020), *VviMYB14* (VIT_205s0049g01020), and *VviMYB30A* (VIT_217s0000g06190). Based on its high transcript levels, *VviMYBPA1* (VIT_215s0046g00170), a homolog of the *A. thaliana* R2R3-MYB genes *TRANSPARENT TESTA 2*/*AtMYB123* (At5g35550) and *AtMYB5* (At3g13540), appears as an important regulator. It is worth noting that another homolog of *AtMYB123* and *AtMYB5*, namely *VviMYBPAR*, also shows high transcript levels from end of August to September. Another R2R3-MYB gene that stands out due to high transcript levels is *VviMYB174* (VIT_218s0001g09850), a homolog of *AtMYB73* (At4g37260) and *AtMYB77* (At3g50060). As expected for organs/tissue not accumulating anthocyanins, *VviMYBA1* (VIT_202s0033g00410), an ortholog of the *A. thaliana* R2R3-MYB gene *PRODUCTION OF ANTHOCYANIN PIGMENT 1*/*AtMYB75* (At1g56650), is not significantly expressed in young buds.

Several genes encoding TFs of the WRKY type show a substantial increase of transcript levels at the gene expression pattern switch during July (e.g. *VviWRKY20* (VIT_207s0005g02570) or *VviWRKY31* (VIT_210s0003g02810)). Examples for the opposite change, i.e. reduction of transcript levels during July, are *VvWRKY23* (VIT_207s0031g01840) and *VviWRKY41* (VIT_213s0067g03140). High transcript levels that are strongly reduced towards winter were detected for *VviWRKY25* (VIT_208s0058g01390) that is, together with *VviWRKY41*, homologous to *AtWRKY54* (At2g40750), *AtWRKY70* (At3g56400) and *AtWRKY46* (At2g46400). Like some of the *R2R3-MYB* genes, also several *VviWRKY* genes display high transcript accumulation values specifically in November and/or an increase towards winter. These include *VviWRKY16* (VIT_206s0004g07500), *VviWRKY45* (VIT_214s0108g01280) and *VviWRKY33* (VIT_211s0037g00150) that are all homologous to the *A. thaliana WRKY* genes of group I-C including *AtWRKY33* (At2g38470), *AtWRKY58* (At3g01080), and *AtWRKY32* (At4g30935).

### 2.4. Identification of qRT-PCR Reference Genes

The quite long RNA-seq time series from tissue of compound buds, covering changing day length and oscillating weather conditions in the field, allowed the identification of candidate reference genes for quantitative Real-Time PCR (qRT-PCR) experiments. The 20 best candidates for reference genes in qRT-PCR experiments were predicted based on an overall high expression and a low variation in steady-state transcript abundance (File S9). A manual inspection of the functional annotation of these genes supported the quality of this data set since it covers well-known qRT-PCR reference genes like ‘polyubiquitin’, ‘glyceraldehyde-3-phosphate dehydrogenase’, ‘elongation factor Tu’, and ‘actin’ as top candidates. The most promising candidate is *VviUBQ10* (VIT_219s0177g00040) which is homologous to the five *A. thaliana* genes encoding polyubiquitin (At4g05320 and others). The second best candidate is VIT_219s0015g01090, a homolog of *A. thaliana HEAT SHOCK PROTEIN 81.4* (At5g56000).

## 3. Discussion

This time series of 18 RNA-Seq data points throughout the first year of development of *V. vinifera* compound buds allows the investigation of developmental processes at relatively high resolution. A similar time series analyses has been performed previously with Affymetrix arrays to determine gene expression patterns for the cultivar Tempranillo [9]. While the data from Tempranillo that was grown near Madrid cover the time from May (first year) to April (second year) with 8 time points, the focus of the time series presented here was the early bud development in June until November of the first year. Nevertheless, the previously reported expression pattern changes in July and also towards winter (November time point) [9] were essentially matched by our data set.

Our chronologically dense collection of samples allows the detection of small developmental differences between time points, but it is affected by a high variation through individual differences between plants, sampled buds, and varying environmental conditions like temperature and other weather conditions. Harvesting the buds in the field is technically challenging since the buds are small and need to be cut out of the axil between shoot and leave or from hard wood. The sampling has to be performed quickly and there was no time to remove adhering tissue from shoots before freezing the samples in liquid nitrogen. The results presented were derived from data of the year 2016 and from time points for which three independent biological replicates were available. However, there are more data of time points for which individual replicates were lost mainly during the RNA extraction procedures due to the problematic technical properties of the respective samples (File S1). While these time points should not be used for statistical analyses, they still provide additional support for patterns observed or may increase the power of co-expression analyses in the future.

Predicted reference genes for qRT-PCR experiments of bud samples contain commonly used reference genes like GAPDH, actin, polyubiquitin, and elongation factor Tu. Additionally, novel candidate genes were identified which displayed a constant expression level. This aligns well with a previous study, which reported that novel reference genes identified by genome-wide *in silico* analysis outperformed typical reference genes in common wheat [32].

We selected three well-known and in the model plant *A. thaliana* well-characterised TF gene families for more detailed analyses of gene expression patterns. MADS-box genes were selected because of their relation to development, and also because quite some of these genes were analysed in the Tempranillo study of Diaz-Riquelme et al. [9]. R2R3-MYB genes and WRKY genes were selected because of their link to stress responses as well as control of accumulation of specialized metabolites.

The gene *VviSOC1a*, a potential integrator of multiple flowering signals that cumulate in the establishment of inflorescence meristems [33], is expressed in June and expression declines after middle of August, when *VviAP1* and also *VviLFY*/*VviFL* (VIT_217s0000g00150, see File S3; ortholog of *AtLEAFY* [34], At5g61850) show increasing expression levels. This increased expression of potential inflorescence meristem identity genes (*AP1*, *LEAFY*) coincides with proliferation of inflorescence primordia, giving rise to inflorescence branch primordia in the developing compound buds [5,6,35]. Based on the also increasing expression of the dormancy marker gene *VviDRM1*, it can be postulated that other parts of the compound bud are already in July on their way to endodormancy. Also the gene *VviTM8a*, a homolog of *TOMATO MADS 8* that plays a role in tomato flower development [36], displays an interesting expression pattern that hints at inflorescence developmental processes taking place during June and beginning of July.

With respect to the expression of the two R2R3-MYB factors known to control proanthocyanidin (PA) biosynthesis that displayed conspicuous expression patterns, namely *VviMYBPA1* and *VviMYBPAR* [37,38], three prominent potential target genes *VviLAR1* (VIT_201s0011g02960) *VviANS* (or *VviLDOX*, VIT_202s0025g04720) and *VviANR* (VIT_200s0361g00040) show expression patterns expected for targets (File S3), also with a reduction of expression levels towards winter. The three genes encode the enzymes leucoanthocyanidin reductase (LAR), anthocyanidin synthase (ANS, also referred to as LDOX) and anthocyanidin reductase (ANR) which are required for biosynthesis of catechin and epicatechin that are precursors of PAs [38]. This indicates that the compound buds accumulate PAs during summer and fall in preparation for winter. Similarly, it is conceivable that the activation of *VviMYB14* and *VviMYB15* results in the synthesis of stilbenes [39,40], which fits to the activation of three of the *V. vinifera* stilbene synthase genes [41] that are clustered on chromosome 16. While *VviSTS35* (VIT_216s0100g01070) and *VviSTS41* (VIT_216s0100g01130) might be targets of *VviMYB15* based on strong co-expression in November, *VviSTS36* (VIT_216s0100g01100) fits better as a potential target of *VviMYB14*. The analysis of *VviSTS* genes [41] also covered three *VviCHS* genes (*VviCHS3*: VIT_205s0136g00260, *VviCHS1*: VIT_214s0068g00920 and *VviCHS2*: VIT_214s0068g00930). The three *VviCHS* genes are co-expressed with a pattern similar to that of *VviLAR1*, *VviANS* and *VviANR* which fits the substrate requirement for catechin/epicatechin biosynthesis and to *VviMYBPA1* and/or *VviMYBPAR* as potential regulators. The homologs of the highly expressed gene *VviMYB174*, *AtMYB73* and *AtMYB77*, have been implicated in root-related auxin responses [42]. A similar function of *VviMYB174* would fit to its high expression throughout compound but development from June to November.

Of the *VviWRKY* genes that have been implicated in the control of *VviSTS’s*, namely *VviWRKY03*, *VviWRKY24*, *VviWRKY43* and *VviWRKY53* [40], only *VviWRKY03* (VIT_201s0010g03930) displayed expression with a pattern that fits to that of *VviMYB14* and *VviSTS36*, allowing to hypothesize that STS36 expression is under combinatorial control of MYB14/WRKY03 in compound buds. *VviWRKY25*, and to some extent *VviWRKY41* as well, that are both homologous to the *AtWRKY* genes implicated in brassinosteroid-regulated plant growth [43], show expression patterns that fit to growth actions until September that are then abandoned towards November and winter.

To the best of our knowledge, this is the first report of a correlation between HSP gene expression patterns with the environmental temperature in a comprehensive time series of field (vineyard) samples. However, repeated formation of HSPs was previously described in the seeds, seed pods, and flowers of *Medicago sativa* [44]. The occurrence of HSPs at standard (not stressed) growth conditions indicated a potential role in development [44]. The annotation “heat-shock” was initially introduced based on up-regulation of genes in heat stress experiments [11,45]. HSPs were also detected at substantial levels in field-grown *Gossypium hirsutum* under increased temperature and drought stress [46]. Reports from *Oryza sativa* support the stress signal integration function of HSPs [47]. Therefore, we also checked for other stress factors like documented pathogen attack, crop protection treatments, or drought stress, but temperature was the only factor with a substantial correlation. Considering the large number of physiological functions of HSPs, constantly expressed chaperons might also be annotated as HSPs. The observation of constant high expression for a gene annotated as “heat-shock protein” (VIT_219s0015g01090) among the potential reference genes for qRT-PCR supports this hypothesis. This assumption aligns well with previous findings that HSPs can have functions in the integration of stress signals [48]. Since our analysis of HSPs is based on the currently available functional annotation, it is likely that the gene expression correlation of bona fide HSP genes with the temperature profile might be even stronger than described here. Moreover, it is possible that additional factors like an underlying developmental pattern or UV-B exposure have an additional influence on the observed heat-shock gene expression profile.

## 4. Materials and Methods

### 4.1. Biological Material

Buds of consecutive time points within the first year of their developmental cycle were taken from a vineyard of the cultivar ‘Calardis Musqué’ (‘Bacchus Weiss’ x ‘Seyval’), former breeding line GF.GA-47-42 (VIVC variety number 4549; http://www.vivc.de). The plot consists of 1300 vines, planted in 1995, pruned as a single cane Guyot system, and located at the Institute for Grapevine Breeding Geilweilerhof in Siebeldingen (49°13’05.0“N 8°02’45.0”E), about 120 m north of a weather station (https://www.am.rlp.de/Internet/AM/NotesAM.nsf/amwebagrar/). Early in the afternoon on each sampling date of the growing season, bud samples were taken in triplicates. From three different vines four buds each were harvested in a batch and immediately frozen in liquid nitrogen. The buds were taken from the fourth to eight node of the shoots emerging from the middle section of the cane (Figure 4). Vines that appeared to be equal in their overall developmental stage were chosen. Those that showed symptoms of nutrient deficiency or diseases were excluded as a sample source.

**Figure 4:**
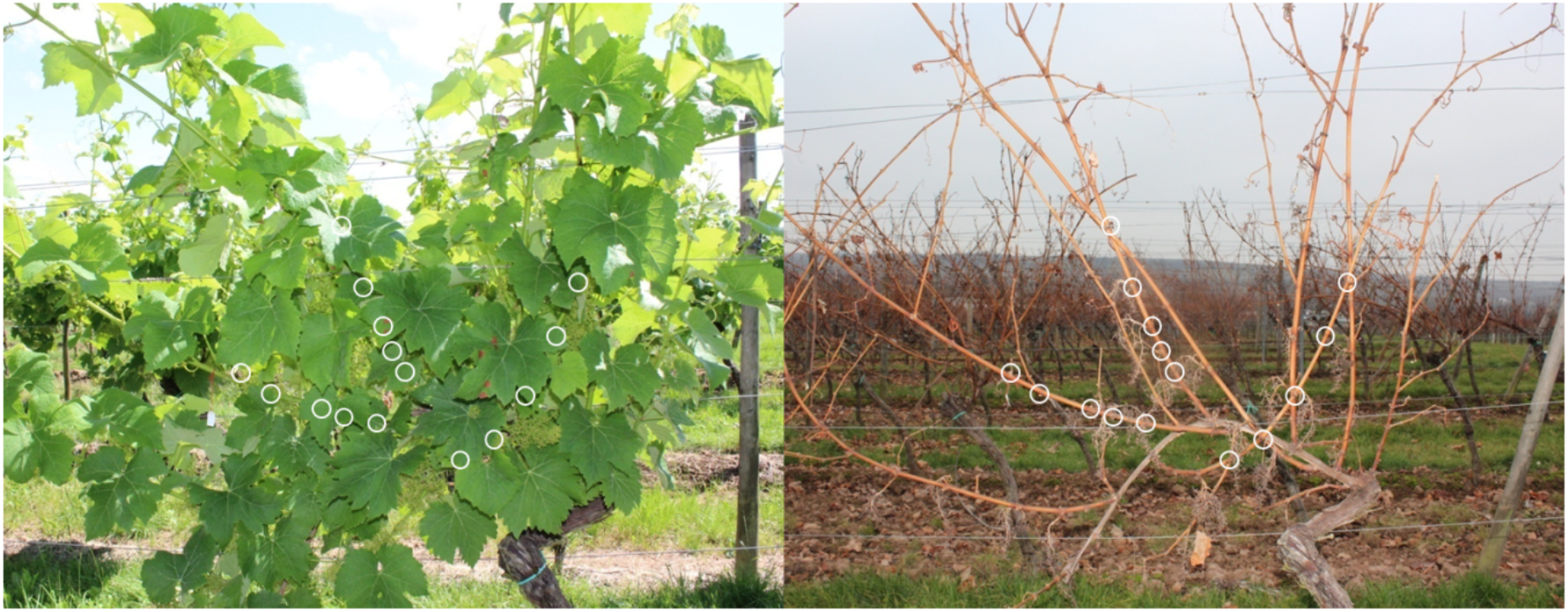
The same vine of ‘Calardis Musqué’ on 1st of July and 16th of December 2016. White circles mark the fourth to eight node of the middle shoots from which the buds were harvested.

### 4.2. RNA Extraction, Library Preparation, and Sequencing

Total RNA was extracted, from four buds each, in triplicate per time point. Up to 100 mg of liquid nitrogen ground tissue was applied to the SpectrumTM Plant Total RNA kit (Sigma-Aldrich, Taufkirchen, Germany) according to the manufacturer’s instructions for protocol B. After on-column DNase treatment with the DNase I Digest Set (Sigma-Aldrich, Taufkirchen, Germany), the RNA was quantified. 500 ng total RNA per sample were used to prepare sequencing libraries according to the Illumina TruSeq RNA Sample Preparation v2 Guide. Purification of the polyA-containing mRNA was performed using two rounds of oligo(dT) oligonucleotides attached to magnetic beads. During the second elution of the polyA+ RNA, the RNA was fragmented and primed for cDNA synthesis. After cDNA synthesis, the DNA fragments were end-repaired and A-tailing was performed. Multiple indexing adapters, specific for each libray and sample, were ligated to the ends of the cDNA fragments and the adapter ligated fragments were enriched by 12 cycles of PCR. After qualification and quantification, the resulting sequencing libraries were equimolarly pooled and sequenced generating 100 nt single-end reads on eight lanes of an Illumina HiSeq1500 flowcell at the Sequencing Core Facility of the Center for Biotechnology (CeBiTec) at Bielefeld University.

### 4.3. Bioinformatic Analysis of RNA-Seq Data

All RNA-Seq read data sets generated were submitted to the ENA (for accession numbers see File S1). Python scripts developed for customized analyses are available at Github: https://github.com/bpucker/vivi-bud-dev. RNA-Seq reads were mapped to the CRIBI2.1 reference genome sequence of PN40024 [49] via STAR v.2.51b [50] with previously optimized parameters including a minimal alignment length cutoff of 90% and a minimal similarity cutoff of 95% of the read length [51]. FeatureCounts [52] was deployed for quantification of steady-state transcript levels at the gene level based on these mappings and the CRIBI2.1 annotation [53]. Previously developed Python scripts [51] were applied to merge the resulting count tables and to calculate counts per million (CPMs) and reads per kb per million mapped reads (RPKMs). We attempted to include *VviFT* (GSVIVT00012870001 in the Vv8x genome sequence) in the analyses of selected target genes, but the corresponding sequence region is not included in the genome sequence version (file Vv12x_CRIBI.fa) on which the CRIBI2.1 annotation is based. To include the three *VviCHS* genes [41], structural gene annotation was optimised for VIT_214s0068g00920 and VIT_214s0068g00930.

Average day temperature values were retrieved from the weather station in the vineyard for the time from June 1st to November 3rd of 2016.

HSP genes in CRIBI2.1 were identified based on the annotation text of homologs in *A. thaliana* by filtering for the strings ‘heat’ and ‘shock’ occurring together in the functional annotation text of the genes. Lowly expressed genes were excluded from downstream analyses by applying a minimal CPM cutoff of 10 (per gene sum over all samples). The Python package matplotlib v2.1.0 [54] was used for visualization of the data.

Members of the transcription factor families WRKY [16], MADS-box [23], and MYB [19,39] were identified based on the published gene family analyses. The *V. vinifera* WRKY gene family has also been characterised by Guo et al. (2014) [55] which, unfortunately, resulted in conflicting gene designations. For consistency with Vannozzi et al. [40] we only used the *VviWRKY* gene designations of Wang et al. (2014). To allow the expression analysis of all previously described MADS-box genes, the CRIBI v2.1 annotation was extended with corresponding gene models using “VIT_230_” as prefix for the additional locus IDs. The Python packages matplotlib v2.1.0 [54] and seaborn v0.8.1 (https://github.com/mwaskom/seaborn) were used for visualization of RPKM values of selected genes in heatmaps.

Candidates for reference genes suitable for qRT-PCR experiments in the future were identified based on our comprehensive set of RNA-Seq samples. First, genes with a substantial expression level defined as sum of all samples greater or equal to 500 [CPM] were selected. Second, these candidate set was filtered for a low variation defined as small standard deviation values across all samples normalized by the median of all values.

## Supplementary Materials

The following files are available online at www.mdpi.com/xxx/s1:

File S1: RNA-Seq sample overview including ENA accessions and number of reads per sample,
File S2: Raw counts of RNA-Seq reads mapped to CRIBI2.1,
File S3: CPMs of RNA-Seq reads mapped to CRIBI2.1,
File S4: RPKMs of RNA-Seq reads mapped to CRIBI2.1,
File S5: IDs of potential heatshock genes in CRIBI2.1,
File S6: Gene expression heatmap of genes encoding MADS-box TFs,
File S7: Gene expression heatmap of genes encoding R2R3-MYB TFs,
File S8: Gene expression heatmap of genes encoding WRKY TFs,
File S9: List of candidates for qRT-PCR reference genes.

## Author Contributions

DH, LH, RT and BW conceived and designed research. BP, AS, SB, and PV conducted the experiments. BP, LH, DH and BW interpreted the data. BP, DH, and BW wrote the manuscript. All authors read and approved the final version of the manuscript.

## Acknowledgments

We are grateful to Willy Keller for excellent technical assistance. In addition, the authors wish to thank all members of the group “Genetics and Genomics of Plants" and the Bioinformatics Resource Facility of the CeBiTec for their excellent assistance and support. We also acknowledge the financial support of the German Research Foundation and the Open Access Publication Fund of Bielefeld University for the article processing charge.

